# Identifying a Vaginal Microbiome-Derived Selective Antibiotic Metabolite via Microbiome Pharmacology Analysis

**DOI:** 10.1101/2025.08.28.672927

**Authors:** Smrutiti Jena, Damilola Lawore, Javier Muñoz Briones, Kenzie Birse, Alana Lamont, Romel D. Mackelprang, Laura Noel-Romas, Michelle Perner, Xuanlin Hou, Elizabeth Irungu, Nelly Mugo, Samantha Knodel, Stephanie Doran Brubaker, Timothy R. Muwonge, Elly Katabira, Sean M Hughes, Fernanda L. Calienes, Raymond Krajci, Rachel Liu, Dalí Nemecio, Florian Hladik, Jairam Lingappa, Adam D Burgener, Alicia R Berard, Leopold N Green, Douglas K. Brubaker

## Abstract

The vaginal microbiome plays a critical role in maintaining immune and epithelial homeostasis in the female reproductive tract. Bacterial Vaginosis (BV) is deleterious to female health, causing the loss of beneficial *Lactobacillus* species, overgrowth of anaerobic taxa, changes in vaginal pH, breakdown of protective mucins and epithelial barriers, and activation of the immune system. Treatment with gel-based antibiotics (Metronidazole or Clindamycin) resolves BV for 85% of patients, but 50% of those cases recur, indicating a need to identify strategies for overcoming antibiotic resistance and achieving a more durable response. Here, we developed a systems biology approach termed *Microbiome Pharmacology Analysis* to characterize the antibiotic potential of vaginal microbes, their metabolites and functions, via computational fusion of human cohort multi-omics and post-drug perturbation transcriptomic profiles. We focused on Clindamycin and Metronidazole as candidate drugs and screened 780 vaginal microbiome-drug mimicry candidates to identify candidate taxa and metabolites with antibiotic potential. We demonstrate experimentally that *Lactobacillus crispatus*-derived Hydroxyisocaproate (HICA) selectively kills *Gardnerella vaginalis* and that HICA enhances epithelial barrier integrity in a human vagina-on-a-chip system. Our work demonstrates the first use of *Pharmacobiome Analysis*, for discovering novel, selective antibiotic metabolites for BV with implications for charting the full pharmacologic potential of the vaginal microbiome.

## INTRODUCTION

The female genital tract is a complex microenvironment comprised of an immune-competent epithelial barrier tissue and microbiome that work in concert to maintain mucosal homeostasis and resist invasion by pathogenic organisms^1^. The interactions between epithelial cells in the female reproductive tract (FRT) are governed by intercellular junctional molecules, including tight junctions that regulate paracellular permeability, as well as adherens and desmosomal proteins, which act as a physical defense against the movement of microorganisms across the epithelial barrier^2^. *Lactobacilli* and their metabolic products may exert direct effects on the mucosa of the FRT, contributing to beneficial outcomes such as improved wound healing, strengthened epithelial barrier integrity^3^, and the maintenance of a non-inflammatory state^4^. The vaginal microbiome is typically comprised of *Lactobacillus* species, including *L. crispatus, L. iners, L jensenii, and L. gasseri*, which exhibit many beneficial properties such as the production of lactic acid, maintenance of low vaginal pH^5^, and formation of protective biofilms^6^. Bacterial vaginosis (BV) is a condition of vaginal dysbiosis characterized by loss of *Lactobacillus* species and overgrowth of facultative anaerobes such as *Gardnerella, Mobiluncus,* and *Prevotella*. This change in vaginal microbiome ecology occurs concomitantly with vaginal epithelial disruption, increased production of inflammatory cytokines^7^, and symptoms including odor, discharge, pain during sexual intercourse or urination, and itchiness^8^.

Metronidazole and Clindamycin are the clinically recommended antibiotics for treating BV, but recurrent and resistant disease remains a significant challenge^9^. Approximately 50% of patients present with recurrence within one year of treatment^10^, with factors such as past BV infection, sexual behaviors, and ethnicity implicated in recurrence^11, 12^. Many alternative strategies are being investigated as BV therapeutics^13^, with a strong emphasis on those that target or use the vaginal microbiome directly as a therapeutic agent, including vaginal microbiome transplant^14^, pH modulation^15–17^, biofilm disruption^18, 19^, and probiotics^20–22^. The lack of effective therapies for BV is a major challenge for female reproductive health as BV associates with and increases susceptibility to many other conditions such as preterm birth^23^ and transmission of sexually transmitted infections^24^. It has also been implicated in pathologies such as infertility^25^, endometriosis^26^ and gynecologic cancers^27, 28^. Since BV impacts as many as 3 million women in United States annually^29^, the discovery of new therapeutic strategies to prevent and treat BV remains a critical need.

In this work, we address the need for efficacious alternatives to antibiotics for treating and preventing BV using a systems biology multi-omics modeling approach that fuses microbiome and pharmacology analysis that we term “*Pharmacobiome Analysis”*^30^. Pharmacobiome Analysis elucidates novel therapeutic functions of microbiome constituents and byproducts by integrating *in vitro* drug screening with human *in vivo* microbiome multi-omics. We compared post-drug treatment transcriptomics data for Clindamycin and Metronidazole from the NIH Common Fund Library of Integrated Network and Cellular Signatures (LINCS) database^31^ with vaginal microbiome-host gene expression signatures derived from secondary analysis of vaginal microbiome multi-omics data from the Partners PrEP cohort^32–34^. The logic behind this approach is that if a microbiome component (e.g., microbe, metabolite, or function) is associated with a gene expression change in the host that is similar to one caused by one of the drugs, that microbiome factor might itself be a beneficial drug. In total, we screened 780 combinations of vaginal microbiome factors and drug candidates and experimentally tested the most promising candidate identified by Pharmacobiome Analysis using bacterial culture metabolomics, bacterial growth and killing assays, and a vaginal epithelial organ-on-chip system.

## RESULTS

### Creation of the vaginal pharmacobiome network resource

Pharmacobiome Analysis begins by extracting (1) drug-gene and (2) vaginal microbiome factor-gene signatures for comparison of candidate microbiome factor-drug mimicry relationships (**Figure 1**).

**Figure 1.**
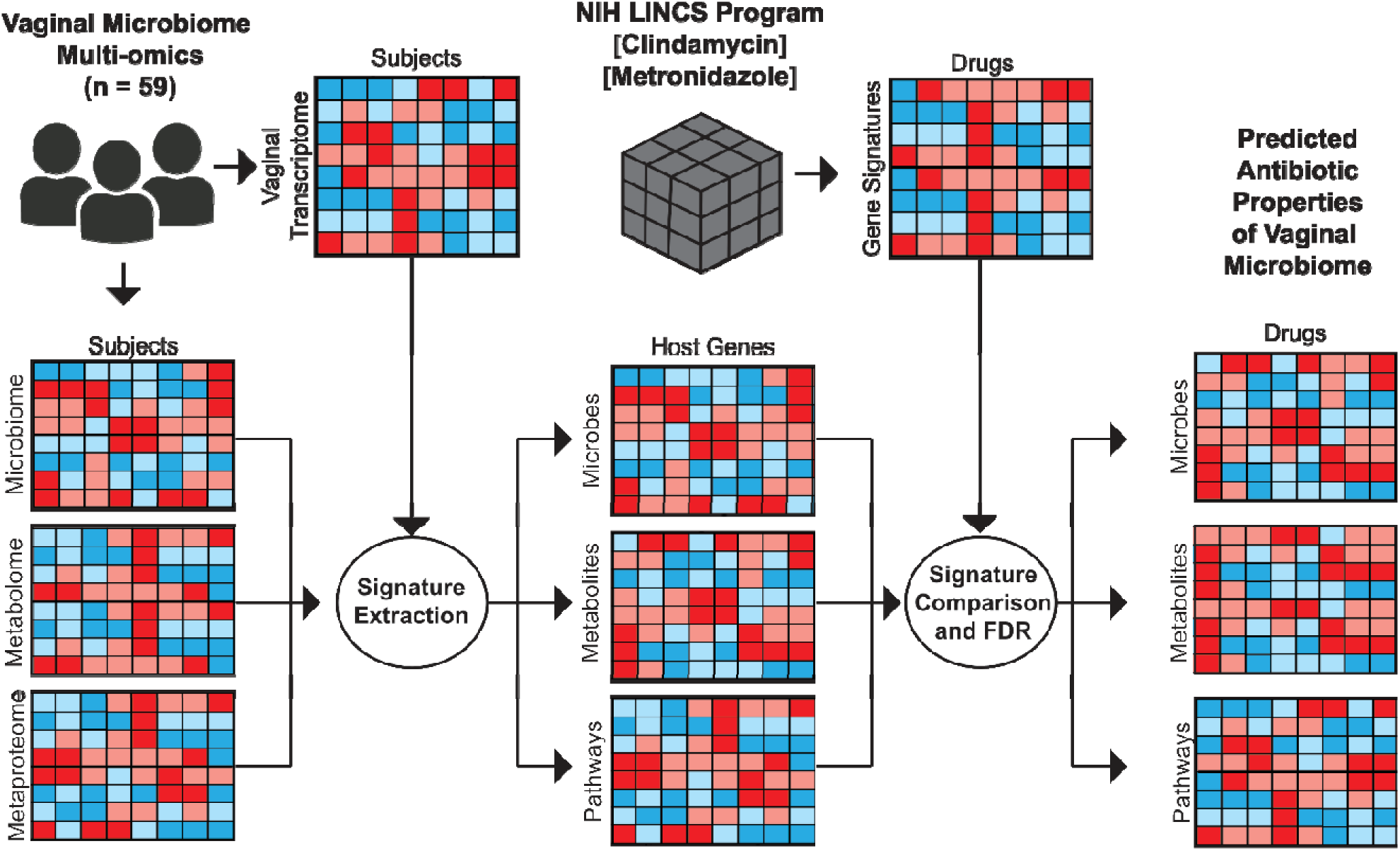
Overview of Vaginal Pharmacobiome Analysis. **(A)** Vaginal microbiome multi-omics data is integrated with matched host transcriptomics via Spearman correlation analysis to generate microbe-gene, metabolite-gene, and bacterial functions-gene lists. Human vaginal microbiome-host gene lists are combined with LINCS characteristic direction signatures via Spearman correlation with False Discovery Rate correction to identify vaginal microbiome-drug mimicry associations.

To extract the drug-gene signatures, post-drug treatment perturbation gene expression data were obtained from the NIH LINCS database at the level of the *Characteristic Direction* (CD) score for the “Chemical Perturbations” category for each drug-gene pair for Metronidazole and Clindamycin^31^. CD profiles aggregate the post-treatment gene expression data across all cell lines, drug doses, and time points to obtain a consensus gene signature for the drug. We chose this approach rather than a specific cell line, treatment concentration, or time point because no vaginal cell lines were included in the LINCS database.

To extract the vaginal microbiome factor-gene signatures, we obtained vaginal microbiome multi-omics data from a subset of HIV-negative participants enrolled in the Partners PrEP clinical trial^32, 34^. We analyzed data from 59 participants that included vaginal microbiome composition, vaginal metabolomics, vaginal metaproteomics inferred via mass spectrometry analysis, and vaginal epithelial transcriptomics data obtained from vaginal biopsies. Spearman correlation analysis was used to calculate a per-gene correlation coefficient in the host vaginal transcriptomics data for each microbe, metabolite, and bacterial function variable in the vaginal microbiome composition, metabolomics, or metaproteomics data. The result was a correlation matrix for each microbiome data type quantifying how the abundance of a microbe, metabolite, or bacterial pathway activity is related to the expression of genes in vaginal epithelium, with the coefficients approximating a magnitude and directional change of the gene in response to the microbiome factors.

Having extracted the sets of drug-gene relationships and vaginal microbiome factor-gene relationships, we next connected them to infer similarities between the vaginal microbiome factors and drugs. Specifically, we generated a matrix of potential vaginal microbiome factor-drug mimicry inferred associations by calculating a Spearman correlation coefficient for each vector of microbiome factor-gene coefficients and drug-CD gene coefficients from LINCS. At this stage, we applied a Benjamini Hochberg FDR correction to the microbiome-drug correlation coefficient p-values to identify statistically significant linkages for downstream visualization and analysis (**Figure 1**). We found 100 metabolites, 165 taxa, and 125 bacterial functions potentially associated with host gene signatures that were also affected by the two antibiotics (2 drugs x 390 microbiome factors = 780 candidate mimicry associations). After filtering out missing values and technical dropouts, our total numbers of metabolites, taxa, and bacterial functions associated with host gene signatures impacted by the two antibiotics were 100, 13, and 32, respectively (**Table S1**).

### Identifying vaginal bacterial abundance and function relationships to antibiotic drugs

Having computed the drug mimicry correlation values for Clindamycin, Metronidazole, and all pairs of vaginal microbes, metabolites, and bacterial pathways, we identified the top candidate microbiome factors likely to mimic the function of these antibiotics (**Figure 2A-C**). We noted that *L. iners, L. crispatus, Gardnerella,* and *Bifidobacterium* mapped to the extremes of the distribution. High positive correlation similarity existed between the host gene signature associated with *Gardnerella* and *Bifidobacterium* and the cell line gene signatures associated with the antibiotics. In contrast, there was a negative correlation similarity for *L. iners* and *L. crispatus* and the antibiotics.

**Figure 2.**
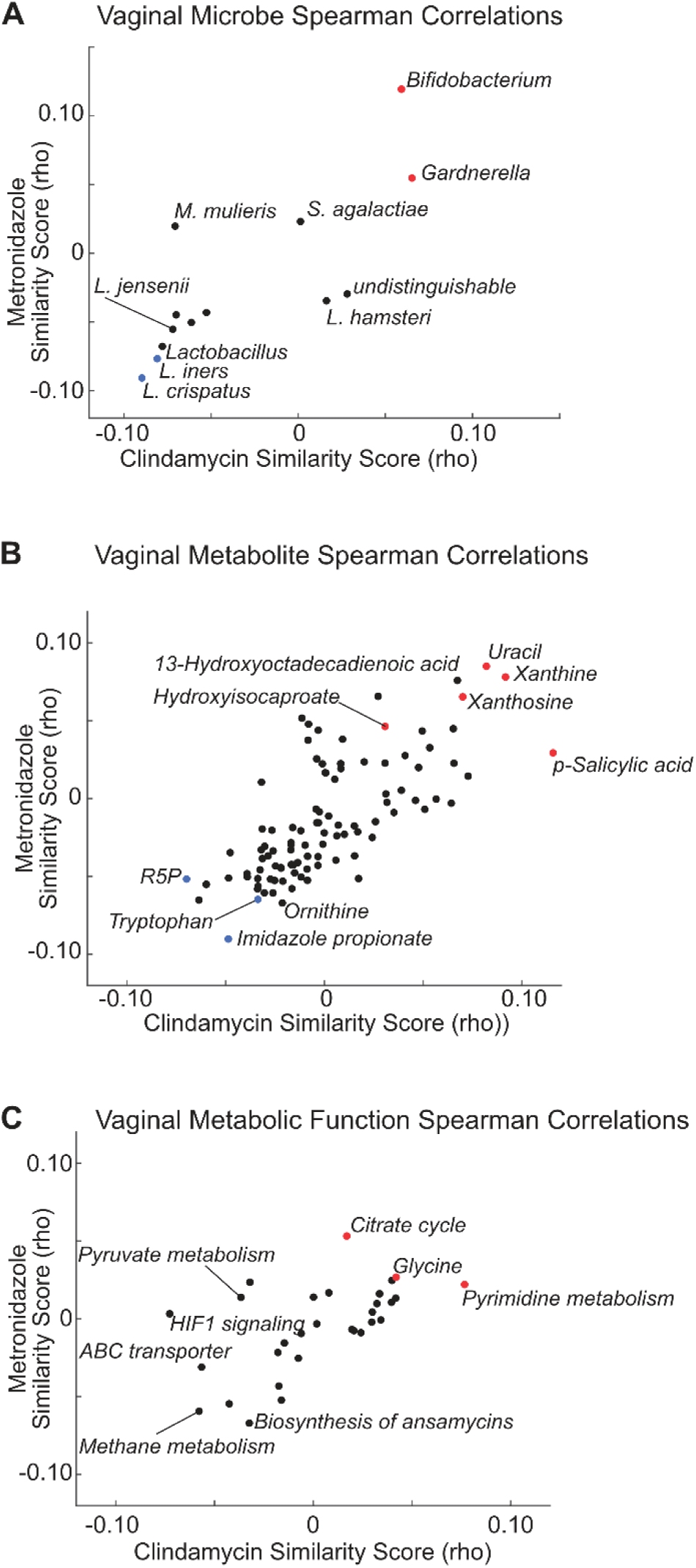
Identification of Vaginal Microbiome Factors with Antibiotic Potential. Shown are correlation similarity scores (Spearman’s rho) for **(A)** relating the correlation coefficients between vaginal microbes and host gene expression to the CD perturbation values of clindamycin and metronidazole; **(B)** relating the correlation coefficients between vaginal metabolites and host gene expression to the CD perturbation values for clindamycin and metronidazole; **(C)** relating the correlation coefficients between vaginal bacterial metabolic functions and host gene expression to the CD perturbation values for clindamycin and metronidazole. See Methods for calculation of correlation similarity scores.

We identified several promising drug candidates among the microbiome metabolites based on strong protective gene correlations with both antibiotics. Specifically, we identified Xanthine, Xanthosine, Uracil, Hydroxyisocaproate (HICA), and p-Salicylic acid as potential antibiotics (**Figure 2B**). The anti-correlated metabolites common to both antibiotics (strongly negative correlation similarity scores) included Tryptophan, Ribose 5 Phosphate (R5P), and Imidazole propionic acid. Among these, Imidazole propionic acid was found to be associated with increased inflammation, activation of mTOR signaling, and BV status^35^. Finally, we identified bacterial functions that, when activated, induce similar host gene signatures to Metronidazole and Clindamycin, including the Citrate cycle, Pyrimidine metabolism, and Glycine signaling (**Figure 2C**).

### Hydroxyisocaproate is a microbiome-derived metabolite selectively antibiotic against BV-associated bacteria

In examining our candidate metabolites from Pharmacobiome Analysis, we noted several positive results already known to have antibiotic functions, including p-Salicylic acid and Uracil. Salicylic acid promotes the antimicrobial effect of Carbapenem against *P. aeruginosa* and has anti-gonococcal activity against azithromycin-resistant *N. gonorrhoeae* with minimal growth inhibition of commensal vaginal bacteria^36, 37^. Uracil, while conjugated with Gentamycin or Ciprofloxacin, increases susceptibility of methicillin-resistant *Staphylococcus aureus* (MRSA) to the antibiotic treatment^38, 39^.

In addition to these metabolites for which antibiotic activities are already known and which can derive from host cell metabolism, we sought to identify novel microbiome-derived molecules that could show antibiotic potential. To identify metabolites of potential vaginal microbiome origin, we performed untargeted metabolomics analysis of conditioned media and matched control media for vaginal strains (*G. vaginalis, L. crispatus, L. iners*) grown in biofilm and suspension conditions (**Figure S1A-B**). Among our top candidate antibiotic metabolites HICA was found to be produced by most vaginal strains, with *L. crispatus* producing the greatest absolute amount on average (suspension: 7.8, biofilm: 13.0) compared to *L. iners* (suspension: 3.6) and *G. vaginalis* (suspension: 5.5, biofilm 1: 6.4, biofilm 2: 0.19) **(Figure S1A)**. The variation in HICA production when scaled to media (**Figure S1B**) suggests that HICA could be produced by the vaginal microbiome, including *G. vaginalis,* but depends on specific nutrient availability.

Next, we performed preliminary experiments to assess the antibiotic potential of HICA by titrating concentrations of 0, 5, 25, and 50 mM cultured with a BV-associated *G. vaginalis* strain for 24 hours (**Figure S1C**). These experiments indicate that 25mM of HICA is enough to confer 73% growth inhibition of *G. vaginalis*. As a comparison and control, we characterized the toxicity of 20mM HICA in 24 hour growth with *L. crispatus* and found only 8% inhibition of growth (**Figure S1D)**.

Having identified 20mM HICA as a dose well tolerated by *L. crispatus* and 25mM as an antibiotic dose for *G. vaginalis*, we harmonized the doses and protocol and performed time course, dose-response, and pH monitoring experiments of cultured *L. crispatus* and *G. vaginalis* to assess the antibiotic potential of HICA at lower doses and earlier time points (**Figure 3**). We found that for all HICA concentrations (5 mM, 10 mM, 20 mM), *L. crispatus* continued to grow in biomass over 24 hours and showed no decrease in viability (**Figure 3A**), but a decrease in culture pH potentially due to lactic acid produced by *L. crispatus* (**Figure 3B**). In contrast to *L. crispatus, G. vaginalis* showed a decrease in viability at all concentrations of HICA, starting as low as 5mM and achieving significant inhibition with 20mM, that caused near total elimination of *G. vaginalis* from culture (**Figure 3C**). We measured the pH of the media for *G. vaginalis* and found no changes in pH during bacterial growth, but that HICA itself reduced the pH of the media in a dose-dependent manner (**Figure 3D**). Therefore, the mechanism by which HICA appears to act as an antibiotic is by modifying environmental pH, a similar manner to which *L. crispatus* maintains vaginal homeostasis *in vivo* and a known mechanism by which the vaginal microbiome resists colonization by pathogens and BV-associated bacteria^40^.

**Figure 3.**
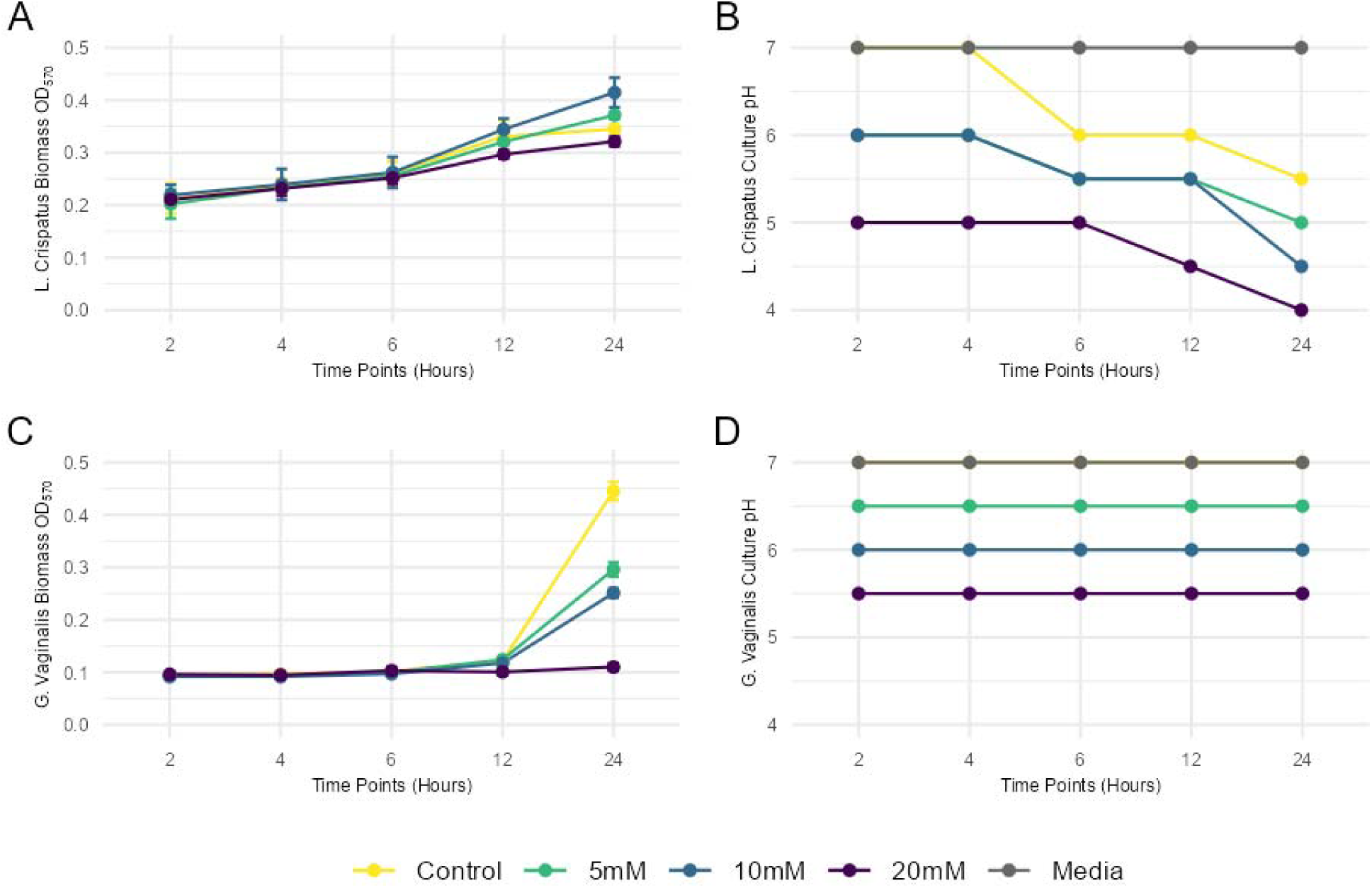
(A) *L. crispatus* growth curve with different concentrations of HICA treatment. (B) Media pH changes at different timepoints of *L. crispatus* growth with HICA treatments (C) *G. vaginalis* growth curve with different concentrations of HICA treatment. (D) Media pH changes at different timepoints of *G. vaginalis* growth with HICA treatments.

### Hydroxyisocaproate enhances epithelial barrier integrity in a vagina-on-a-chip organoid system

Though HICA showed efficacy as a selective vaginal microbiome-derived antibiotic in bacterial culture, the impact of potential therapeutic metabolites on host physiology is equally important to consider for translational viability. To assess the suitability of HICA for translation as a therapeutic for BV in patients from the perspective of impact on vaginal barrier integrity, we implemented the Emulate organ-on-a-chip system to create a vagina-chip (as previously described^41^). The Emulate microfluidic vagina-chip contains stratified multilayered epithelium and primary human stromal fibroblasts, mimicking the underlying connective tissue of vaginal mucosa. The epithelial layers are connected by tight junctions, adherens junctions and desmosomes, and fibroblasts provide mechanical stability and elasticity to the tissue^1^. This configuration maintains a physiological oxygen gradient besides enabling simulation of physiological parameters including nutrient flow rate and hormone fluctuations^41^.

Once the vagina chip system was established, we introduced 20 mM of HICA. Post 48hr incubation, we assessed for the impact of HICA on cellular integrity and barrier permeability (**Figure 4**). After 48 hours, the HICA-treated system showed increased cellular density and better formation of an epithelial layer compared to the untreated condition, indicating HICA enhanced the quality of the epithelial barrier of the vagina-chip (**Figure 4A**). We also assessed the effect of HICA on inter-epithelial barrier integrity and permeability by Fluorescein isothiocyanate–dextran dye permeability assay. As with the visual inspection via Brightfield microscope showing a more intact epithelial barrier, the permeability experiment quantified an increase in epithelial barrier integrity and reduced permeability (**Figure 4B**). These data together show that the introduction of HICA improved overall vaginal epithelial barrier integrity potentially as the same mechanism as lactic acid by regulating pH^42, 43^. Together with the bacterial culture experiments, this demonstrates HICA is a potentially translatable selective antibiotic, that inhibits *G. vaginalis* growth with minimal impact on *L. crispatus* growth, and eubiotic metabolite for BV treatment.

**Figure 4.**
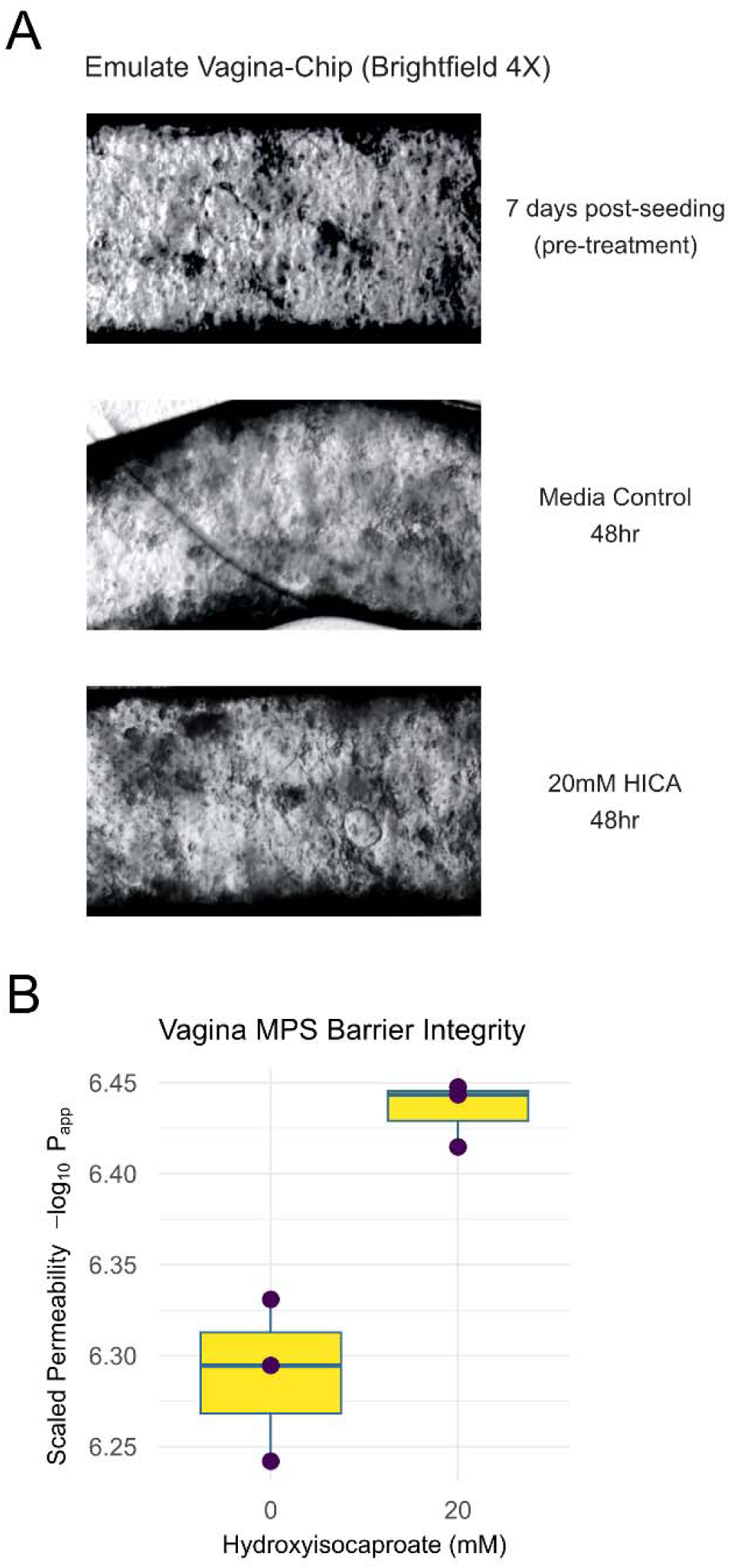
(A) Brightfield microscope image of vagina-chip system treated with 20mM of HICA versus pre-treatment and media control. (**B**) Apparent permeability of the vagina-chip epithelial layer after 48hours of HICA treatment shows increased barrier integrity (-log10(apparent permeability)).

## DISCUSSION

Here, we leverage a combination of vaginal microbiome multi-omics analysis and drug screening data to create a systems biology platform termed *Pharmacobiome Analysis*. Using this platform, we identify a candidate BV therapeutic and confirm its activity in the lab. Our novel Pharmacobiome Analysis pinpoints the therapeutic functions of microbial constituents and byproducts. We screened 780 microbiome-drug mimicry candidates to arrive at a ranked list of microbes, metabolites, and bacterial functions with antibiotic potential. Metabolomics analysis of cultured *L. iners, L. crispatus,* and *G. vaginalis* allowed us to distinguish between host and microbiome-derived metabolites and prioritize potential therapeutic metabolites by their ability to be produced by the beneficial strain *L. crispatus*. Finally, screening of HICA, our candidate antimicrobial compound, in bacterial cultures demonstrated its potential as a *G. vaginalis*-selective antibiotic, and analysis of HICA effects on a vaginal epithelial organ-on-chip showed this metabolite enhances vaginal barrier integrity. Taken together, we present a novel therapeutic strategy with HICA that has the potential to ameliorate multiple hallmarks of BV, including the overgrowth of anaerobic bacteria, support of a *Lactobacillus* community, and enhancement of vaginal epithelial barrier integrity.

HICA is a metabolite in fermented foods, including cheeses, soy sauce, wines, and kimchi^44^. It is derived from the branched-chain amino acid leucine by hydroxyisocaproate dehydrogenases (HicDs)^44^. Evidence suggests that the fermentation process’s specific lactic acid bacterial composition influences HICA production, effectively inhibiting the growth of food spoilage and foodborne pathogens. Its mechanism of action Includes penetrating bacterial cell membranes, leading to depolarization, membrane disruption, and cell death^45^. HICA exhibits broad-spectrum activity against clinical pathogens, including *Candida, Aspergillus, Staphylococcus*, and *Fusobacterium* species, and periodontal pathogens such as *Porphyromonas gingivalis* and *Fusobacterium nucleatum*^46^. Moreover, HICA has shown efficacy against multi-drug-resistant strains, including *Pseudomonas aeruginosa*, while being well tolerated in host tissue, even illustrating delayed muscle soreness^47^ and repair^48^. To our knowledge, this study represents the first exploration of HICA’s application in the vaginal context, highlighting its potential as a novel antimicrobial agent in this niche.

Other potential antibiotic metabolites implicated in our study include Xanthine, Xanthosine, Uracil, and p-Salicylic acid. Our cultured bacterial metabolomics analysis indicated that NAD+, gluconate, glucose-6-phosphate, harmane and dihydroferulic acid can be produced by *Lactobacillus* strains as well. Of these, gluconate and harmane have the potential to enhance the effect of other antibiotics^49, 50^ with gluconate being able to promote wound healing^51^, further supporting Pharmacobiome Analysis’s ability to identify microbiome-derived metabolites’ with clinically relevant functions. Many of these metabolites have not been tested in the vaginal tract or as therapeutics for BV, making them potential candidates for novel therapeutics along with HICA.

Though our overall hypothesis that the magnitude of the similarity score is predictive of drug mimicry seems to hold for metabolites, in the taxa-based Pharmacobiome analysis (**Fig 2A**), we observed that *Gardnerella* was positively associated with the antibiotics scores and *Lactobacilli* were negatively associated. This result is opposite of what one might expect given the positive health characteristics associated with a *Lactobacillus* dominant vaginal community. Our interpretation of this is that since the antibiotics reduce the abundance of *Gardnerella* the genes positively associated with *Gardnerella* abundance would decrease and those genes negatively associated would increase. The effect of this would be to invert the expected correlation and produce the apparent, but incorrect inference that higher levels of *Gardnerella* would induce host gene responses similar to antibiotic treatment. This highlights that accounting for the effect of a drug on the microbiome’s ecology and abundance of taxa is important in taxa-based Pharmacobiome inferences.

Our approach takes advantage of Characteristic Direction-averaged gene expression profiles from LINCS to define the vaginal pharmacobiome network, integrating data from various cell lines, time points, and drug concentrations into a single profile. The absence of vaginal cell lines in LINCS is a limitation, since gene expression profiles derived from Clindamycin and Metronidazole-treated vaginal epithelial cells would be more representative of the site where these drugs act. Additionally, optimizing dosing concentrations and time points in drug perturbation transcriptomics experiments could yield higher-fidelity data for vaginal microbiome-based drug discovery. Despite these limitations, the averaged gene expression profiles enabled experimentally testable predictions and the validation of antibiotic-candidate metabolites. The biological context provided by the Partners PrEP multi-omics data likely contributed to overcoming the absence of vaginal cell lines in LINCS. Despite these limitations, we show that the averaged gene profiles of the CD led to experimentally testable predictions and validated antibiotic candidate metabolites. Further refinement of cell type-specific drug perturbation transcriptomics would enhance our analysis and represent a promising future research direction. The logic behind Pharmacobiome Analysis is similar to the Connectivity Map project^31^ where perturbation gene expression signatures of drugs were used to identify repurposing candidates. While the Connectivity Map approach has limitations such as a potential high false positive rate^52^, we control for that here through stringent FDR levels, prioritization of microbiome-derived molecules through *in vitro* metabolomics analysis, and rigorous testing of therapeutic effects experimentally.

This study makes a complementary contribution to the body of work harnessing systems biology approaches to understand the vaginal microbiome and its role in BV^34, 53, 54^. Indeed, addressing the many critical unmet needs in female reproductive health will require an array of methods tuned to address specific challenges. We envision Pharmacobiome Analysis as a broadly applicable, encompassing framework with the potential to accelerate microbiome-based therapeutic discovery and enable precise characterization of the therapeutic potential of all microbiomes. Further in depth molecular and pathway level studies will facilitate insight into the mechanism of the eubiotic function of the bacteria derived compounds/metabolites advancing towards their therapeutic implementation.

## METHODS

### Human Vaginal Microbiome Multi-Omics Data

Vaginal microbiome multi-omics data from a subset of 405 HIV-negative women enrolled in the Partners PrEP (Pre-Exposure Prophylaxis) study^32^. The proteomics, metaproteomics, and metabolomics data were previously generated (as described^35^), and we used data from a subset of 90 participants who were profiled for transcriptomics data from vaginal biopsies as well as the above measures. Our analysis here focused on the subset of matched multi-omic samples from Group 2 participants, with our largest subset of matched data covering 59 patients.

### Drug Perturbation Transcriptomics Data

Post Clindamycin and Metronidazole treatment gene expression profiles were obtained from the Library of Integrated Network and Cellular Signatures (LINCS) Chemical Perturbation SigComm database^31^. Data were obtained at the level of *Characteristic Direction* (CD) level which summarizes the net perturbation gene expression profile of a drug across multiple cell lines, concentrations, and time points (CD perturbation value). The rationale for using the averaged CD perturbation value rather than cell line-specific responses for the drugs is because there were no vaginal epithelial cell lines in LINCS, thereby making the consensus CD signature the best of the available signature options.

### Integration of Drug and Microbiome Data

Microbiota composition, metaproteomics, and metabolomics data from the Partners PrEP cohort subset were analyzed via Spearman correlation analysis with the vaginal epithelial transcriptomics data to identify a gene signature for each microbe, microbial pathway, and metabolite. The magnitude and direction of the correlation value is treated like a hypothesized regulatory relationship. The vector of gene-based Spearman correlation values for each microbiome factor was then correlated (also Spearman) against the CD perturbation values for Clindamycin and Metronidazole from LINCS to generate a correlation similarity score for each drug-to-microbiome factor pairing. These correlation similarity scores for microbiome factor-drug pairings were subjected to multiple hypothesis correction via Benjamini Hochberg, preserving the multiple comparison correction for the stage of the workflow where the results of the hypothesis test are interpreted.

### Vaginal Bacterial Metabolomics

The cultured bacterial metabolomics data used in this study was performed as described and data are deposited in Metabolomics Workbench: http://dx.doi.org/10.21228/M82M8R. In brief, *L. crispatus* ATCC 33820, *G. vaginalis* ATCC 14018, and *L. iners* ATCC 55195 were cultured for the metabolomics study. For *L. crispatus*, De Man–Rogosa–Sharpe (MRS) media was used for suspension cultures. NYC III media was used for growing *G. vaginalis* suspension. *L. iners* suspension culture was grown in BHI^55^. *L. crispatus* biofilm was grown in MRS media. *G. vaginalis* biofilm was grown in NYCIII and Brain Heart Infusion media supplemented with 2% gelatin, 0.5% yeast extract and 0.1% soluble starch (sBHI) separately to attain two types of biofilms based on substrate used. All cultures were incubated at 37^0^C, 5% CO_2_, 24hr shaking for suspension and 48hr static for biofilm. Post incubation, cultures were centrifuged at 13000 x g for 10mins and then sterile filtered using 0.2μm syringe filter. Cell free supernatants were sent in triplicates to Metabolon, NC, USA, for non-targeted metabolomics^34^.

### Vaginal Bacterial Growth Inhibition Assay

*Lactobacillus crispatus* strain ATCC 33820 and *Gardnerella vaginalis* strain ATCC 14018 were grown in MRS and TSB+5% sheep blood media respectively. Single colony was picked and inoculated in appropriate broth media and cultured with shaking for 24 hrs. in 5% CO_2_ at 37^0^C until 0.4 at OD_570_nm for the MIC experiment of selected metabolites. MIC experiment was performed on 96 well plates (Corning, 3370) with 1:100 culture to media ratio and 5mM, 10mM and 20mM concentrations of L-alpha-Hydroxyisocaproate (Thermo scientific, Cat: 204810050). Temporal growth/inhibition of *L. crispatus* and *G. vaginalis*, under the effect of HICA was measured using Biotek synergy H1 microplate reader at OD_570_. The experiment was repeated thrice with triplicates each time. The effect of 20mM, 30mM and 40mM D-Ribose-5-phosphate disodium salt hydrate (Thermo scientific, cat. No. AAJ6681503) was also screened (data not shown).

### Vagina-on-chip culture

Dual channel microfluidic system vagina-on-chip devices used for this study were from Emulate Inc. (Boston, MA). The system was established using a previously published method^41^. Briefly, the apical channel with 1 mm W × 1 mm H was seeded with Primary human vaginal epithelial cells (Lifeline Cell Technology) and the basal channel having 1 mm W × 0.2 mm H seeded with Primary human uterine fibroblasts (ScienCell Research Laboratories, cat. no. 7040). Cocktails of Collagen IV (30 μg/mL) from human placenta (Sigma, cat. no. C7521) with collagen I (200 μg/mL) from rat tail (Corning, cat. no. 354236) and collagen I (200 μg/mL) with poly-L-lysine (15 μg/mL) (ScienCell Research Laboratories, cat. no. 2301) were used for apical and basal chamber ECM coating 2-3 hours prior to introducing cells respectively. Seeding density was 1 × 10^6^ cells/mL for fibroblasts and 3 × 10^6^ cells/mL for epithelial cells. The chips were incubated at 37°C with 5% CO_2_ with continuous media flow. Using Zoe fluidic system (Emulate), the apical chamber was perfused with cervical epithelial cell basal medium (ATCC, cat. No. PCS-480-032) at a flow rate of 20 μL/h and basal chamber using fibroblast growth medium (ScienCell Research Laboratories, cat. no. 70940) at 40 μL/h flow rate. After 4 days, the basal media was replaced with in-house differentiation medium for the basal chamber, composition as described elsewhere^41^.

### Metabolite treatments

The well-established and differentiated vagina-on-chips were treated after 8 days with metabolites for 48 hours at 37^0^C and 5% CO_2_ with unchanged flow rates. HICA and D-Ribose-5-phosphate disodium salt hydrate were added to the apical chamber media at concentrations 20mM and 10mM respectively. We used 9 vagina-on-chips (triplicate for each condition including no treatment control) for this experiment.

### Permeability/barrier integrity assay

Fluorescein isothiocyanate–dextran (Sigma, cat. No. FD4-100MG) was used with the apical chamber media at a concentration of 100 μg/mL. After 24 hours, the flow through from the basal and apical chamber outlet was collected and the fluorescence intensity was measured at an excitation wavelength of 485nm and emission at 528nm using plate reader (BioTek, Synergy H1). Apparent permeability (P_app_) was calculated using the following equation:

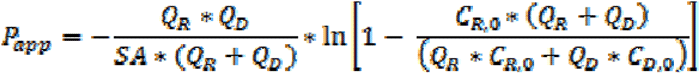

where P_app_ is the apparent permeability in units of cm/s, SA is the surface area of the co-culture channel (0.17cm2), Q_R_ & Q_D_ are the fluid flow rates in the receiving and dosing channels, respectively, in units of cm^3^/s, and C_R,0_ & C_D,0_ are the recovered concentrations in the receiving and dosing channels, respectively, in any consistent units (Emulate, barrier function analysis protocol).

## Data and Code Availability

The vaginal microbiome multi-omics data used in this study is available in its source publication^34^. The LINC CD signatures can be obtained from the SigComm LINCS database^31^. This paper does not report custom computing code.

## ACKNOWLEDGEMENTS

The authors would like to thank the Partners PrEP-BV Study Team and the participants whose data was used in this study. We are grateful for the opportunity to work with the team to advance other areas of gynecologic health.

## AUTHOR CONTRIBUTIONS

Conceptualization: SJ & DKB; Methodology, DL, SJ, DKB; Organ-on-chip: SJ, LNG; Visualization: SJ, DKB.; Writing – original draft, SJ & DKB.; Writing – review & editing: DL, JMB, SDB, ARB, KB, AL, RDM, LNR, MP, XH, EI, NM, SK, TRM, EK, SMH, CL, FLC, RK, RL, DN, JMB, CC, FH, JL, LNG, DKB.

---

**Figure S1.**
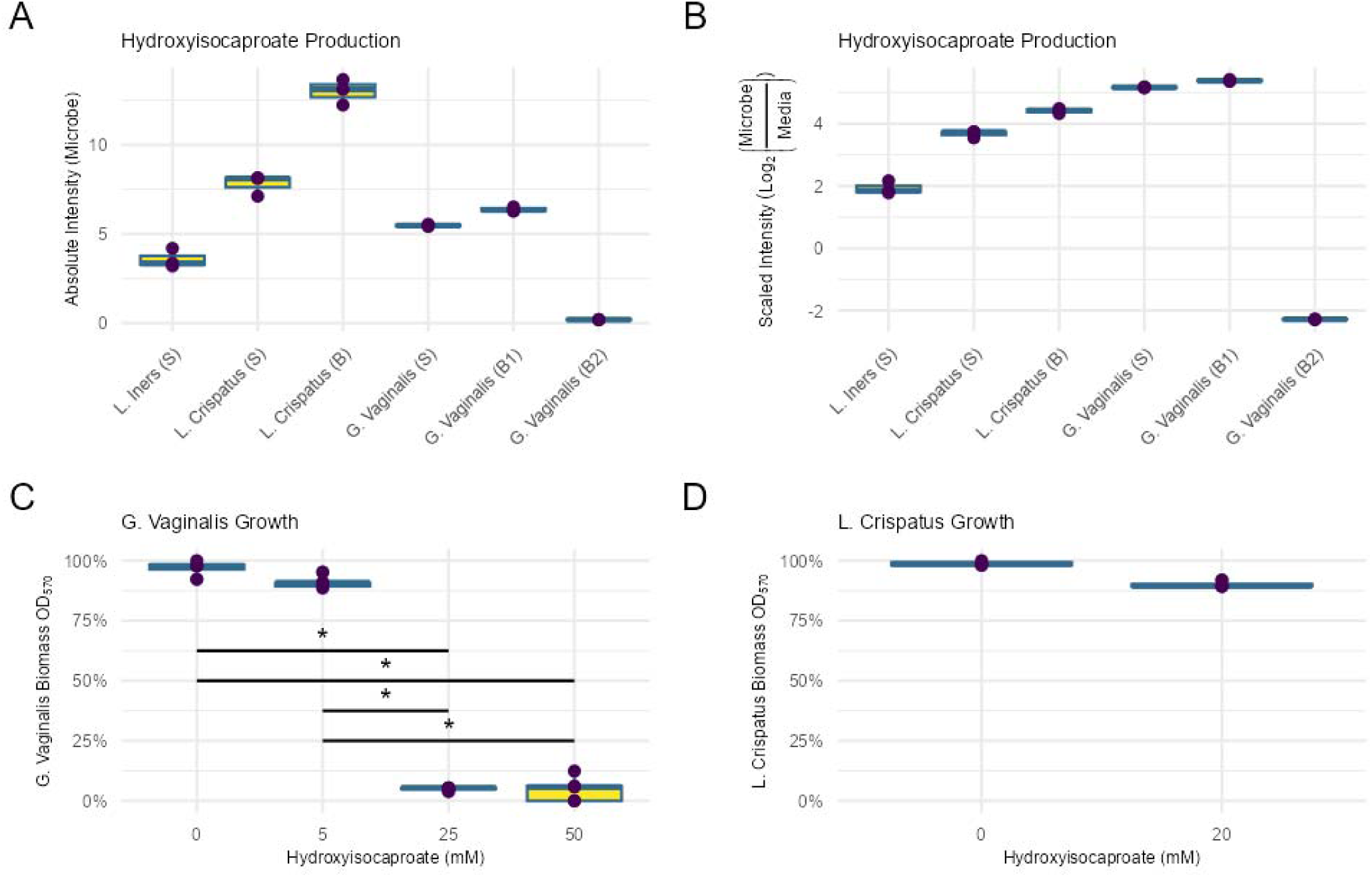
Functional characterization of the antibiotic properties of Hydroxyisocaproate (HICA) **(A)** HICA production by isolated, cultured type strains of vaginal bacteria in suspension (S) and biofilm (B) conditions. *G. vaginalis* biofilms were grown in NYCIII (B1) or supplemented BHI (B2) media. **(B)** HICA production scaled to media **(C)** *G. vaginalis* growth inhibition by HICA treatment. **(D)** *L. crispatus* growth inhibition by HICA treatment.

